# Dark diversity reveals assembly constraints across trophic levels in large-scale grassland restoration

**DOI:** 10.64898/2026.10.06.756980

**Authors:** Qingqing Chen, Laura Forgione, Daniel Slodowicz, Gino Enz, Michael Wyss, Raphaël Arlettaz, Jean-Yves Humbert

## Abstract

1. Grassland biodiversity restoration efforts have largely focused on re-establishing diverse plant communities, with less attention paid to higher trophic levels such as invertebrates.
2. We established a controlled, field-scale restoration experiment across twelve lowland grassland sites in Switzerland in close collaboration with farmers. We manipulated seed addition (hay transfer, directly harvested seeds, or multiplied seed mixtures) and soil disturbance (shallow or deep tillage), with treatments applied to meadows covering, on average, 0.5 ha. Four to six years after restoration, we quantified the components of species richness change (lost, persisting, gained from site surroundings, and gained from donors), and estimated dark diversity (species that are absent from a location but could potentially occur there) for plants, Auchenorrhyncha and Orthoptera.
3. Compared to plants, invertebrates exhibited greater species turnover, with higher proportions of species lost and gained from the surroundings, but much lower contributions from the donor meadows. However, species persistence was high across taxa. Restoration operations strongly influenced components of species richness change in plants but had weaker effects on invertebrates. In particular, multiplied seed mixtures + deep tillage had the highest proportion of plant species not transferred, suggesting that many species in the seed mixture were poorly suited to local conditions. In both plants and Auchenorrhyncha, components of species richness change were strongly mediated by site characteristics, with initial species richness in both receiver and donor emerging as the strongest predictors.
4. Within each taxonomic group, 41% of species not transferred from the donor belonged to the dark diversity, suggesting substantial unrealized restoration potential. Restoration operations impacted dark diversity for plants and Auchenorrhyncha but not for Orthoptera. In plants, dark diversity was substantially lower under multiplied seed mixtures + deep tillage than under other treatments. The probability of a species belonging to dark diversity was associated with specific leaf area in plants and initial relative abundance in Auchenorrhyncha.
5. Synthesis and applications. Components of species richness change were influenced by both restoration operations and site characteristics, with their relative importance varying among taxa, underscoring the need for multi-taxon restoration strategies.

## Introduction

The semi-natural grasslands that historically characterized the cultural landscapes of the Old world have undergone dramatic declines in extent and quality, leading to dramatic losses of plant and invertebrate species richness and shifts in species composition (Bardgett et al., 2021; Gibbs & Salmon, 2015; Suttie et al., 2005). Ecological restoration is widely promoted as a key strategy to reverse these negative trends (Buisson et al. 2022; Dudley et al. 2020). However, restoration efforts in grasslands typically emphasize re-establishing diverse plant communities, but often neglect other trophic levels such as invertebrates (Clark et al. 2007; Török et al. 2024). Invertebrates perform vital functions within agroecosystems, such as pollination, decomposition, nutrient cycling, and food provision for higher trophic levels (Belovsky & Slade, 2000; Garibaldi et al., 2013). Because vegetation composition, structure, and functional trait diversity can shape the assembly of associated herbivore communities, restoring plant diversity is likely to trigger cascading effects across all trophic levels up to the final consumers (Scherber et al. 2010). Seed addition via hay transfer or sowing seed mixtures are methods commonly used for restoring plant diversity in intensively used, species-poor grasslands or arable lands. Numerous studies have shown that such seed addition methods can effectively restore plant species richness (reviewed by Slodowicz et al., 2023). In general, invertebrate restoration positively correlates with plant restoration (Alignan et al. 2018; Alison et al. 2017; Neff et al. 2020; Woodcock et al. 2010). Small-scale experiments have established that several invertebrate taxa can be successfully transferred from a species-rich donor meadow to a receiver meadow via hay transfer (Stöckli et al. 2021; Wagner 2004). Yet, empirical evidence for cross-trophic responses towards effective large-scale realistic restoration practices remains limited.

Partitioning species richness changes into species lost, species gained, and species persisting can provide a more nuanced understanding of biodiversity dynamics in grassland restoration operations. Net changes in richness may mask contrasting underlying processes, whereby similar richness outcomes can arise either from high turnover or strong species retention. Previous grassland restoration studies have merely focused on the number or proportion of species that were successfully transferred from seed mixtures (e.g., Forgione et al., 2026; Freitag et al., 2021; Laschke et al., 2026; Orrock et al., 2023). Expanding that approach by quantifying the probability of not transferred species to dark species (absent from a given location but that could potentially occur there) could further improve biodiversity restoration operations in grasslands (Chollet et al. 2025; Moeslund et al. 2017). For instance, seed mixtures used for restoration can be designed so as to include a larger fraction of those species being more likely to belong to the dark diversity pool. Finally, disentangling the environmental co-factors underlying grassland restoration success will be instrumental in developing more robust methodological options to enhance biodiversity and improve restoration success in grasslands.

Restoration outcomes may be mediated by local community properties (e.g. aboveground biomass and species richness), environmental conditions (e.g. soil nitrogen – hereafter soil N; precipitation), landscape context (e.g. area devoted to restoration and quality of the surrounding matrix), and management history (e.g. years under extensive management) (Freitag et al. 2021; Grman et al. 2010; Orrock et al. 2023; Pywell et al. 2003; Rotchés-Ribalta et al. 2018; Woodcock et al. 2010). In any restoration operations of species-poor grasslands, initial species richness is likely to greatly impact restoration trajectories and success. Sites with a higher initial species richness may lose more species over time, while gaining less from surroundings due to saturation effects and niche occupancy (Cornell, 2013). Similarly, sites with a smaller difference in species richness relative to their donor meadows may have less potential for species gains from the donor. In addition, species’ initial abundance may play an important role in restoration success: rare species are more likely to be lost and less likely to be transferred from donors to receivers because their higher demographic vulnerability and reduced establishment likelihood (Pimm et al., 1988). Despite these well-established theoretical expectations, their implications for biodiversity restoration in grasslands have yet to be empirically tested. Functional traits can also mediate restoration trajectories and outcomes by shaping species establishment, resource use strategies and dispersal dynamics (Freitag et al., 2021; Neff et al., 2020). For example, Belinchón et al., (2020) found that grassland plant species with low dispersal capacity have a higher probability of belonging to dark species. Yet, how these factors jointly impact components of species richness change in restoration operations have, to the best of our knowledge, not been evaluated experimentally for multiple trophic levels.

In fact, despite extensive research, major uncertainties remain regarding the effectiveness of restoration practices at spatial scales relevant to farming management and conservation planning. This is because much of the extant experimental evidence stems from small experimental plots, often without proper spatial replication, which hampers drawing general conclusions (Clark et al. 2007; Freitag et al. 2021; Klaus et al., 2020; Orrock et al. 2023; Pywell et al. 2007). Addressing this knowledge gap is essential for designing restoration strategies that are both ecologically robust, applicable in practice and readily accepted by farmers.

We therefore established a large-scale restoration experiment designed to enhance plant and invertebrate diversity in relative species-poor meadows situated in the Swiss lowlands. We tested the effects of different restoration operations combining three seed-addition methods and two soil-preparation techniques, on components of species richness change and dark diversity among grassland plant and invertebrate communities. Previous experiments have suggested that combining seed addition and soil disturbance is essential to increase restoration success of plants (Bischoff et al. 2018; Freitag et al. 2021), but it is unknown what degree of soil disturbance is ideal. We monitored changes in plants, Auchenorrhyncha, and Orthoptera four, five and six years after the onset of restoration, respectively. We assessed how the different restoration operations impact processes of biodiversity change and how local community properties, environmental conditions, landscape context, and management history mediate these processes. We first hypothesized that directly harvested seeds from nearby donor meadows would be most effective in restoring plant species richness due to local co-evolution with environmental conditions (Zuppinger-Dingley et al., 2014), whereas fresh hay transfer would perform best for restoring the invertebrate community. This is because arthropods can either be directly transferred while attached to the freshly mown vegetation, or because their locally growing host plants are transferred (Alison et al., 2017; Stöckli et al., 2021; Wagner, 2004). Second, we predicted that change in species richness is driven by different processes in plants and invertebrates, with more species gained from the donor in plants than in the invertebrates. Finally, we expected that processes of biodiversity change be shaped by various site characteristics, notably that initial species richness as well as species abundance and dispersal-related traits would exert strong effects.

## Methods

### Study sites

The experiment was conducted in the Swiss lowlands located between the Alps and the Jura mountain ranges. Land-use in this region is generally homogeneous and dominated by high-intensity agriculture (Zingg et al., 2024). In 2018, 60 extensively managed meadows were selected at 12 study sites, with five meadows per site, with sites being >10 km apart and situated between 420 and 760 m above sea level (Fig. S1). These meadows had not been fertilized since at least 2013. Mean annual ambient temperature ranged from 10–13°C and precipitation from 886–1118 mm between 2018 and 2023 (MeteoSwiss, 2025).

### Experimental design

All five meadows of a given study site were located within a 5×5 km square, with a minimum distance of 340 m between two adjacent meadows to ensure sample independence. The size of the study meadows ranged from 0.2–1 ha, with a mean of 0.5 ha. Within each study site, an ancient extensively managed, biodiverse grassland was chosen as the site-specific donor meadow. Donor meadows harboured, on average, 47%, 32% and 29% more species of plants, Auchenorrhyncha, and Orthoptera, respectively, than the receiver meadows (see Table S1 for the total number of species, the number of shared species between donor and receiver meadows, and the number of unique species). Four experimental restoration operations and a control were randomly assigned to the meadows within a study site. No experimental intervention was performed in the control, i.e. neither soil disturbance nor seed addition. The four restoration operations tested were: 1) Hay transfer + shallow tillage; 2) Hay transfer + deep tillage; 3) Direct seeds + deep tillage; 4) Multiplied seeds + deep tillage. Hay transfer consisted of spreading freshly harvested hay from a donor meadow onto a receiver meadow on the same day in June 2019, when the majority of seeds had reached maturity, at a 1:1 area ratio. Direct seeding consisted of sowing seeds collected with a seed harvester (eBeetle; developed by HoloSem; Oberwil-Lieli, Switzerland; https://www.holosem.ch/ebeetle/angebot/) from a donor meadow, mostly in June 2018. However, additional seeds had also been hand-collected in spring and summer 2018 from the same donor meadow so as to include early and late flowering species. The seeds collected with the eBeetle were all sown by hand on the receiver meadows in May 2019 (ca 1.2 g seeds m^-2^). One of the meadows subjected to the direct seeds + deep tillage treatment had to be discarded from the study because extreme rainfall washed the seeds away at the beginning of the experiment. Multiplied seeds refer to a regional native seed mixture produced by multiplication via culture by a commercial company (UFA Salvia CH-G, https://www.ufasamen.ch). This seed mixture comprises 38 plant species with genuine Swiss ecotypes and is the main seed mix used by Swiss farmers to re-instate flower-rich grasslands on arable land. These commercial seeds were mechanically sown onto the receiver meadows in May 2019. Shallow tillage was performed with a rotary harrowing machine a few days before hay transfer and again on the very day of hay transfer. Deep tillage, i.e. ploughing, was performed 2–3 months before seed addition, with raking operations carried out every 4–6 weeks until sowing.

The experiment was initially established to promote plant indicator species, which are used as a criterion for result-based agri-environmental payment schemes in Switzerland. Farmers receive financial rewards when their meadows contain ≥ six indicator species within a 3-m-radius plot. At the beginning of the experiment, these extensively managed meadows generally contained fewer than six indicator species. By 2023, four years after active restoration, 90% of the study meadows contained more than six indicator species, thus meeting the eligibility criterion for larger subsidies (see details in Forgione et al., 2026). The present study used that same experimental set up to investigate the processes underlying changes in plant and invertebrate species richness and dark diversity.

### Vegetation survey

Vegetation surveys were conducted in May–June one year before restoration (2018) and four years after restoration (2023). In each meadow, the vegetation was recorded in two permanent rectangular plots of 2 × 4 m, laid in a perpendicular position (see Fig. 3 in Forgione et al. 2026). These two relevé plots had initially been randomly placed in any given meadow. Within each plot, all herbaceous vascular plant species were recorded and their percentage cover estimated visually.

**Table. 1.**
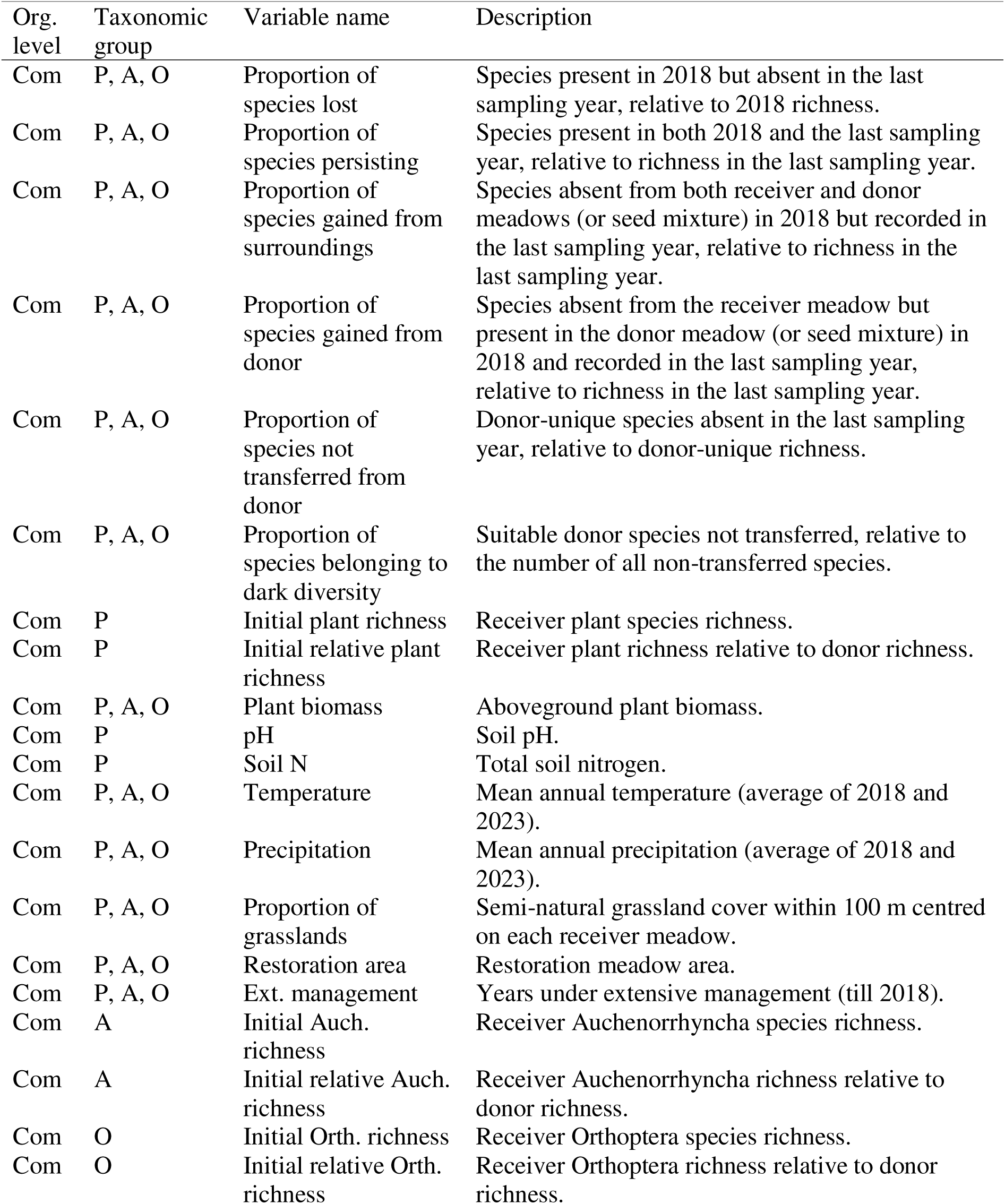
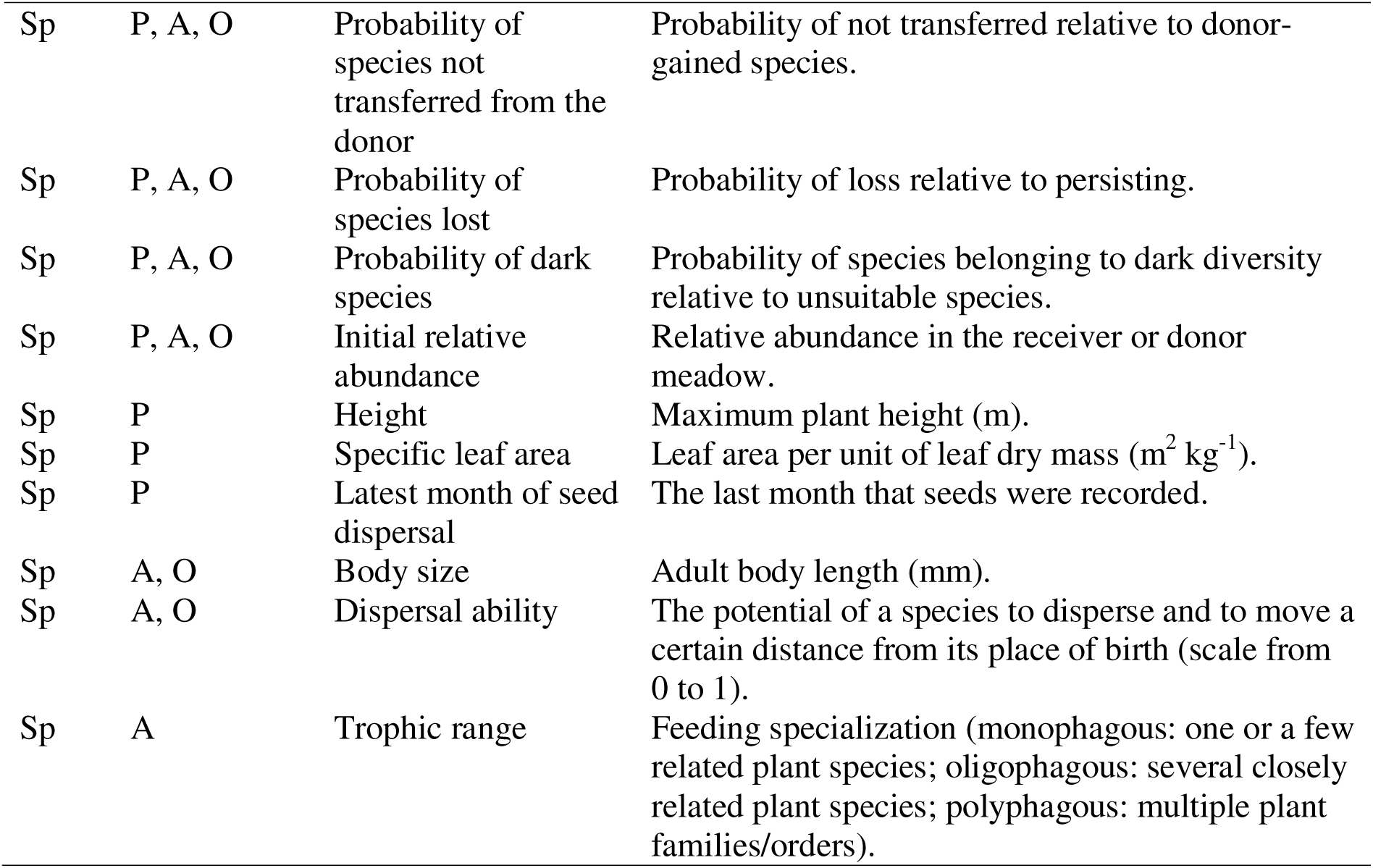
Summary of all variables investigated, with name and description. Abbreviations for the organizational (org.) level: Com = community; Sp = species. Abbreviations for the taxonomical groups: P = plants; A = Auchenorrhyncha; and O = Orthoptera. All site characteristics were measured in 2018 before restoration unless otherwise stated.

### Auchenorrhyncha survey

Auchenorrhyncha were collected in 2018 and 2024 using a suction sampler (Stihl SH86; Stihl, Waiblingen, Germany). The device has an air flow rate of 770 m^3^ h^-1^ and an air suction velocity of 85 m s^-1^. Each sample was vacuumed from inside a circular metallic tube enclosure of 0.51 m diameter (surface area 0.2 m^2^) that prevented arthropods from escaping (Buri et al. 2016). Vacuuming duration was at least 20 s per sample. A virtual cross with 6 m long perpendicular branches was randomly placed between the two vegetation plots. Four samples were collected during a field survey, one at the two extremities of each branch, thus covering a total sampled area of 0.8 m^2^. Samples were collected twice a year: before (end of May – mid-June) and after mowing (beginning of July). They were kept in plastic bags and stored in a deep freezer (-20 °C) until laboratory analyses. Individuals were counted, while adults were identified to species level.

### Orthoptera survey

Orthoptera were sampled in 2018 and 2025 during two annual sampling sessions, one in July, one in August. Sampling was carried out on sunny days between 9 am and 6 pm. Orthoptera species richness and abundance were assessed using a biocenometer, a bottomless cylindrical enclosure made of a rigid net with a sampling area of 1 m² (Buri et al., 2013). In each meadow, 16 biocenometer samples were taken, with a minimum distance of 10 m between sampling points. Adult individuals were identified to species level.

### Functional traits

#### Plants

We opted for height, specific leaf area, and seed dry mass, which have been shown to mediate plant restoration success (Freitag et al. 2021; Neff et al. 2020; Pywell et al. 2003). Additionally, we extracted phenological traits depicting the season of seed maturity, which coincided with the latest month of seed dispersal. Species-level trait data were primarily extracted from TRY database (version 6), supplemented with other published sources (Chen et al. 2024; Kattge et al. 2020). Whenever possible, we used trait data collected in Europe. Yet, for species lacking European information, we used the global average. Overall, these traits could be allocated to 92 % of the 218 species recorded in receiver and donor meadows in 2018.

#### Auchenorrhyncha

We selected body size, dispersal ability, and trophic range for Auchenorrhyncha as these traits have been commonly used and shown to mediate restoration success of invertebrates (Neff et al., 2020; Woodcock et al., 2010). We extracted these trait values primarily from a compiled database, which focuses on northwestern European species (Logghe et al. 2025). This database includes 78% body size, 78% dispersal ability, and 83% trophic range of the 58 species of Auchenorrhyncha recorded in our receiver and donor meadows, respectively, in 2018. Body size refers to the mean length of adults, averaged across males and females. The dispersal ability, estimated by experts, was expressed as the relative potential natal dispersal distance, scaled from 0.1 to 1. Trophic specialization includes 1) monophagous (Auchenorrhyncha feeding on a single or very few related plant species); 2) oligophagous (feeding on several closely related species, typically within the same taxonomic family or genus); 3) polyphagous (feeding on species from multiple families or orders). The remaining 22% of missing data for body size information was obtained from Nickel et al. (2009), while 22% of missing data for dispersal ability was estimated by Dr. Verena Rösch (in litt.). Finally, the remaining 17% of the trophic specialization was extracted from Enz et al. (2021).

#### Orthoptera

We extracted species-specific body size and mobility indices primarily from Logghe et al. (2025), a database comprising information on ca 80% of the 15 Orthoptera species found in our receiver and donor meadows in 2018. Body size values for the remaining 20% of species were obtained from https://www.orthoptera.ch, while mobility indices were estimated by the senior author of this paper, who has extensive experience with this taxon.

### Site characteristics

We distinguished four classes of site characteristics that typically contribute to grassland biodiversity restoration success (Freitag et al. 2021; Orrock et al. 2023): 1) local community properties (aboveground biomass, initial species richness), 2) environmental conditions (soil pH, soil N content, precipitation, ambient temperature), 3) landscape context (e.g. size of the restoration area, proportion of grassland), and 4) management history (e.g. years under extensive management). Aboveground biomass and initial species richness were derived from our surveys conducted in 2018, prior to restoration. Soil pH and soil N were measured in 2018 at each study site using standardized methods. Soil pH was measured potentiometrically in a soil–deionized water suspension using a calibrated pH electrode. Total soil nitrogen (N) concentration was determined by dry combustion using an CNS-Analyzer (vario EL cube by elementar; Langenselbold, Germany). Annual precipitation and ambient temperature data were extracted for each study meadow from the nearest MetoSwiss weather station (distance ranging from 3’624 to 13’622 m) and averaged across the 2018–2023 period. The size of the grassland area being restored was measured for each meadow. The variable “years under extensive management” specified the total duration of the extensive management to which any given meadow was subjected prior to 2018. The proportion of grassland was calculated as the relative cover of semi-natural grasslands within a 100-m-radius area centred on each receiver meadow (see Table 1 for more details).

### Analysis

We partitioned the changes observed in species richness – from the last sampling year (2023 for plants, 2024 for Auchenorrhyncha, and 2025 for Orthoptera) to that of the starting year (2018) – into species lost, species persisting, species gained from the surroundings, species gained from the donor, as well as species not transferred from the donor. For plants, species richness was set as the number of species recorded in the two permanent relevé plots per meadow. For invertebrates, it was calculated as the cumulative number of adults recorded across all samples and sampling sessions within a year. We calculated components of species richness change (species lost, species persisting, species gained from the surroundings, species gained from donor) based on proportional change (see Table 1 for details of quantification of each variable). The proportion of species lost and persisting are relative to the proportion of the total number of species observed in 2018, i.e. before restoration. The proportion of species gained either from a site’s surroundings or from a donor are relative to their proportions in the last sampling year. The sum of the proportions of species persisting, of species gained from a site’s surroundings and from a donor amounted to 100%. The proportion of species not transferred from a donor was relative to the number of unique species, defined as species initially recorded in the donor meadow or seed mixture but absent from the receiver meadows in 2018 (Bischoff et al. 2018; Freitag et al. 2021). The proportions of unique species ranged from 55%–79%, 49%–62%, and 29%–40% for plants, Auchenorrhyncha, and Orthoptera, respectively (Table S1).

We quantified the dark species – corresponding to the probability of a species being ecologically suitable to a receiver meadow but absent from it – by examining, within a site, species co-occurrences across treatments (meadow sites) in 2018 and in the last sampling year, using the hypergeometric method implemented in the R package ‘DarkDiv’ (Carmona & Partel, 2020). This method considers how species *i* that are absent from a meadow in the last sampling year, while being present in other nearby meadows, co-occur with species *j* in that same meadow in any sampling year. We finally quantified dark diversity for any given meadow as the sum of the probabilities of all locally occurring dark species.

We tested the effects of treatments on each component of species richness change using mixed-effect models. Region was included as a random variable, treatment as a fixed effect, while the various components of species richness change were the response variables. For proportional changes, we used the R function glmmTMB from the R package glmmTMB (Brooks et al., 2025) with a beta distribution and a logit link function. For components of species richness change with fewer than 3% zero values, we applied the transformation (y(n-1)+0.5)/n) to satisfy the requirements of the beta distribution, in which *y* is the response value, *n* is the sample size. For components of species richness change with more than 10% of 0 or 1 values (i.e. species gained from the donor for Auchenorrhyncha, species not transferred from the donor for Auchenorrhyncha, species lost for Orthoptera), we restricted analyses to observations lying within the interval (0, 1). This approach was chosen for simplicity and interpretability, allowing the use of standard beta regression without introducing the additional assumptions required for zero–one-inflated models. Sensitivity analyses indicated that alternative treatments of boundary values (e.g. zero–one-inflated models) would not materially affect the results. Statistical analyses for components of species richness with ≤5% of valid observations within (0, 1) (i.e., for Orthoptera, species gained from the donor and species not transferred from the donor) were not conducted due to insufficient variation for reliable model estimation and lack of statistical power, which would have obliterated proper inference. We used the function “emmeans” from the R package “emmeans” (Lenth, 2025) to extract the effect size and used the function “multcomp” from the R package “multcomp” (Hothorn et al., 2025) to perform multiple comparisons between treatments, adjusted using the “Tukey” adjustment method.

We examined whether and how local community properties, environmental conditions, landscape context, and management history impact the different components of species richness change. For Auchenorrhyncha and Orthoptera, we only included variables that can directly impact components of richness change (i.e. Soil pH and Soil N were not included). We used initial species richness to predict the proportions of species lost, persisting, and gained from the surroundings, as all these processes are tightly linked to the receiver community. Relative (to the donor) initial species richness was used to predict species gained from the donor, species not transferred, and dark diversity, as these processes are strongly linked to both receiver and donor communities. We evaluated the relative effects of local community properties, environmental conditions, landscape context, and management history as drivers of each component of species richness change using generalized linear mixed models fitted with glmmTMB. Predictor variables included treatment, initial species richness or initial relative species richness, and all other site characteristics, while region was included as a random effect. All predictor variables were standardized (mean = 0, SD = 1) to facilitate comparison of effect sizes, which represent the change in the logit-scale response per one-standard-deviation change in each predictor. Furthermore, we investigated whether and how changes in the components of species richness in Auchenorrhyncha and Orthoptera were related to changes in plant diversity metrics over time (between 2018 and 2023). Plant diversity metrics were quantified using species richness, Shannon diversity (to account for relative species abundance), and Bray–Curtis dissimilarity (to capture changes in species composition). In the models, the predictor variables included treatment and changes in each plant diversity metric over time, while region was included as a random effect. These results are presented in Fig. S4–S5.

Finally, we investigated whether and how species relative abundance and functional traits impact the probability of species lost (relative to species persisting), the probability of species not transferred (relative to species transfer from the donor), and dark species. For plants, species abundance refers to their relative cover at any given study site in 2018, except for multiplied seeds + deep tillage treatment, where species abundance refers to the relative biomass of the specific seeds with respect to the total biomass of the mixture. For Auchenorrhyncha and Orthoptera, species abundance refers to the number of individuals. We used their initial relative abundance in the receiver meadow to predict probability of species lostlost, while initial relative abundance in the donor served to predict the probability of both species not being transferred and dark diversity. We investigated the relative effects of initial species abundance and functional traits on the probabilities of species lost, species not transferred, and occurrence as dark species using generalized linear mixed models fitted with glmmTMB. Predictor variables included treatment, initial relative abundance, and functional traits, while region was included as a random effect. Similarly, all predictor variables were standardized (mean = 0, SD = 1) to facilitate comparison of effect sizes. All analyses were carried out using (R Core Team, 2025).

## Results

### Plants

Averaged across treatments, the proportion of species lost in receiver meadows was moderate, ranging from 18% to 36% among the 12 sites. In contrast, we observed a high proportion of species persisting, accounting for 49–75% species in the final sampling year. Moreover, 18–25% of species in the final sampling year originated from the surrounding landscape, 21–31% had their origin in the donor (Fig. 2). We found that 52–66% of the unique species were not transferred. Moreover, 16–52% of the not transferred plant species belonged to dark diversity, which corresponded to 3.2–6.9 species in our study meadows (Fig. S3).

**Fig. 1.**
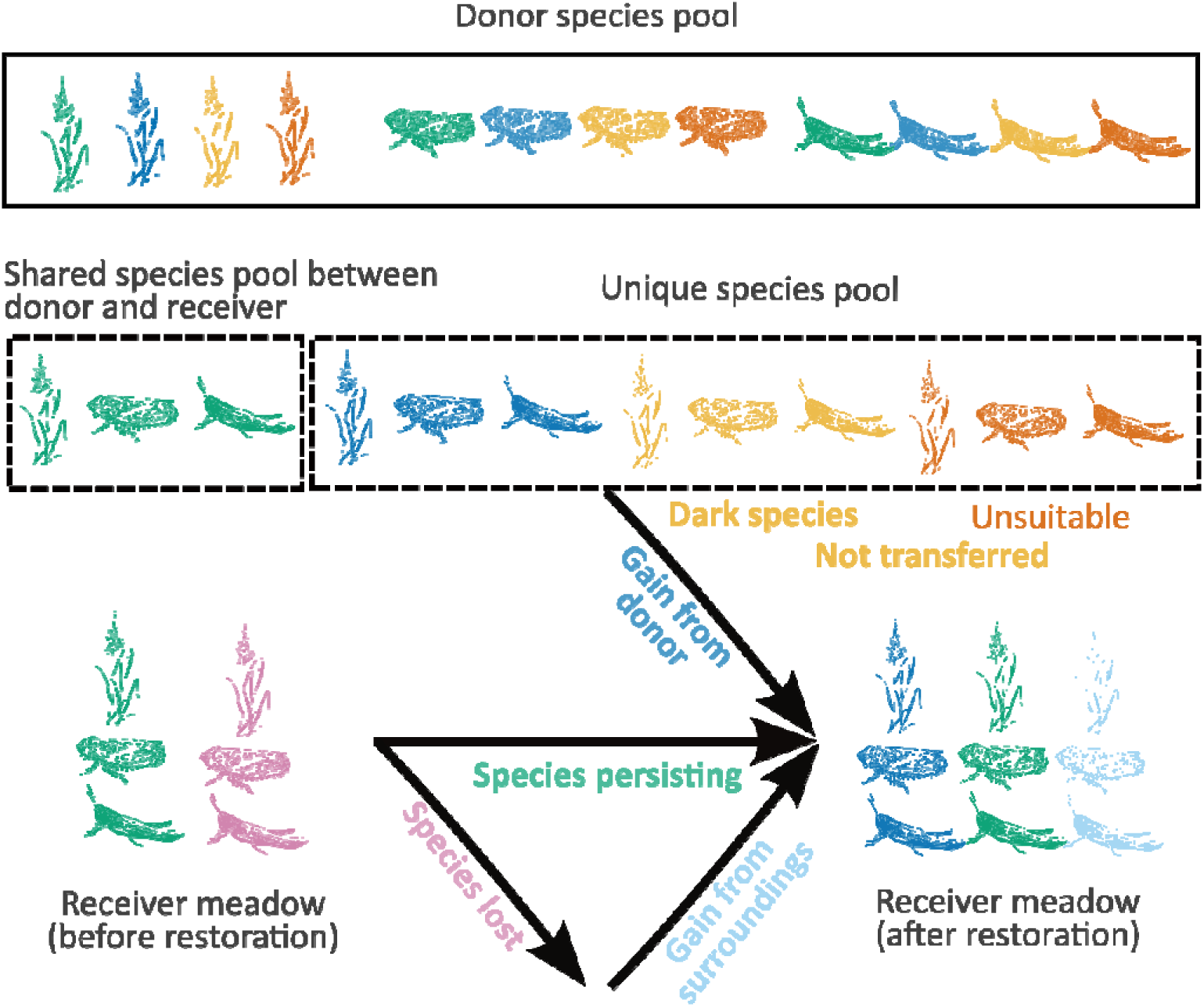
Diagram showing the various components of change in species richness within a grassland restoration framework. Colours symbolize different species of plants, Auchenorrhyncha, and Orthoptera. See Table 1 for a description of the different components/variables.

**Fig. 2.**
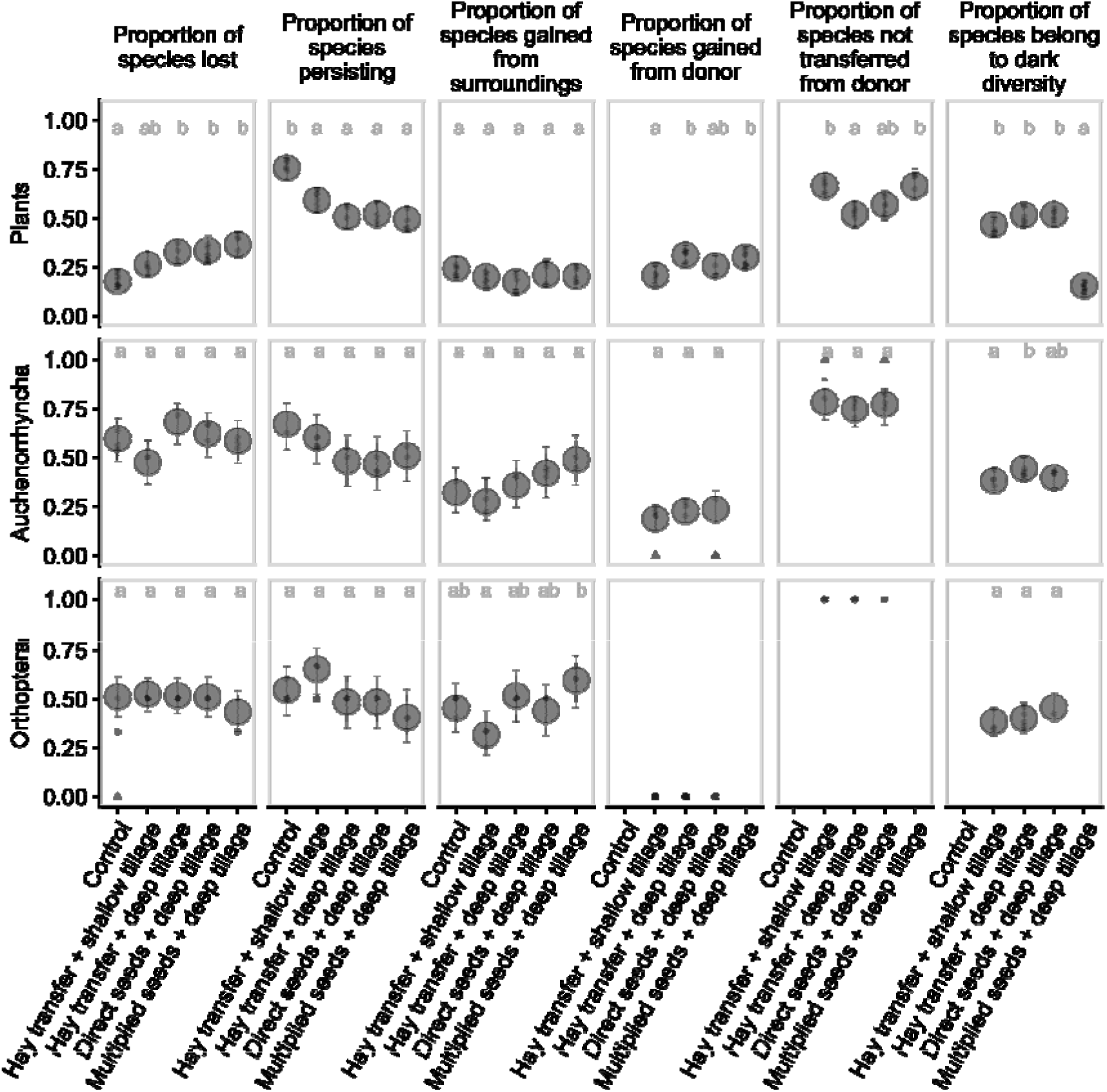
Effects of experimental restoration treatments on components of change in species richness based on proportional change for plants, Auchenorrhyncha, and Orthoptera (components defined in Fig. 1; variable descriptions in Table 1). Small dots show meadow level raw data; triangles indicate observations excluded from the models (see Material and Methods for details). Large dots and related error bars represent estimated means and 95% confidence intervals, respectively. Different letters indicate significant differences at α = 0.05, with Tukey-adjusted multiple comparisons. Fig. S2 shows the effects of the various restoration operations on species richness before and at the last sampling years for the three taxonomic groups; Fig. S3 shows the effects of the restoration operations on the components of change in species richness calculated using absolute changes.

Restoration operations did not change the overall species richness relative to control (Fig. S2), but altered the processes driving species richness change, such as increasing the proportion of species lost while reducing the proportion of species persisting. Among treatments, multiplied seeds + deep tillage resulted in the highest proportion of species lost and the lowest proportion of species persisting. The proportion of species gained from the donor was highest under multiplied seeds + deep tillage and hay transfer + deep tillage, with both treatments exceeding that observed under hay transfer + shallow tillage. Conversely, the proportions of species not transferred from the donor were higher under hay transfer + shallow tillage and multiplied seeds + deep tillage than under hay transfer + deep tillage. Finally, multiplied seeds + deep tillage resulted in the lowest proportion of species belonging to the dark diversity (Fig. 2; Fig. S3; Table S1).

Components of plant species richness change varied substantially and were strongly influenced by multiple site characteristics, particularly initial species richness and soil pH (Fig. 3). Initial species richness was a major determinant of community dynamics: species-rich sites showed higher proportions of both species persisting and species lost, but fewer species gained from the surroundings. Similarly, sites with higher relative species richness compared with donor communities showed reduced contributions of donor-derived species. Soil pH emerged as an important driver of species turnover, being associated with greater proportions of species lost, donor-derived gains, and dark diversity, but lower persistence and fewer species not transferred from donors. Soil nitrogen was positively related to species persistence. Landscape and climatic factors further influenced community assembly, with greater grassland cover reducing species loss and dark diversity; warmer regions showed higher dark diversity than wetter regions. Finally, longer histories of extensive management were associated with reduced species loss, suggesting a legacy effect of past land use on present-day species richness change.

**Fig. 3.**
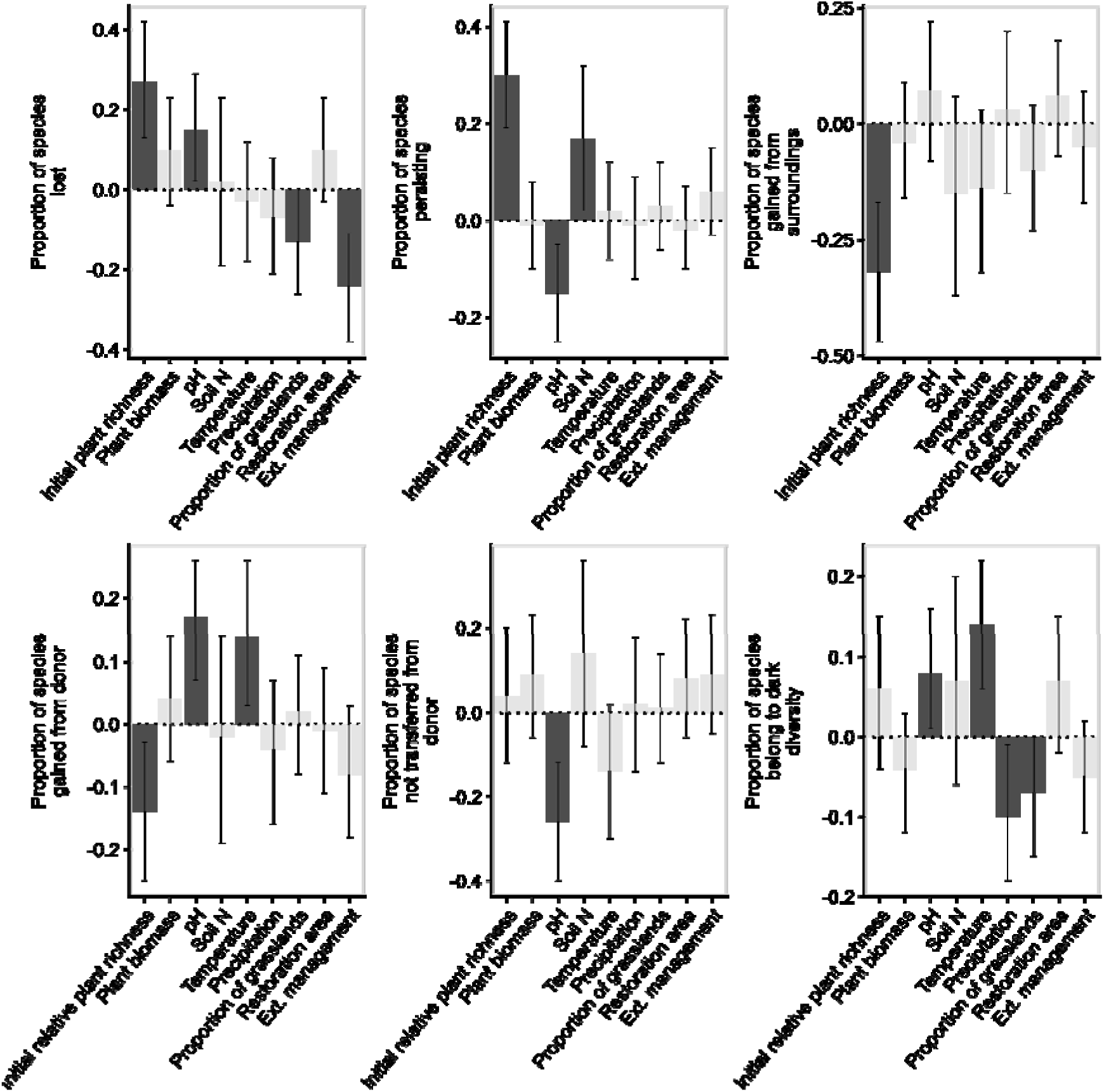
Standardized regression coefficients from multiple regression models for the effects of community properties, environmental conditions, landscape context, and management history on various components of the change in plant species richness based on proportional changes. See Table 1 for the definitions of response and predictor variables. Bars and error bars represent standardized effect sizes and 95% confidence intervals. Dark shading indicates statistically significant effects (α = 0.05), whereas light shading denotes non-significant effects.

At the species level, initial relative abundance rather than functional traits played an important role in mediating probability of species lost, species not transferred from donor, and dark diversity. Plant species with higher initial relative abundance in receiver meadows showed a lower probability of being lost. Similarly, species with higher relative abundance in donor showed a lower probability of not being transferred. We found that plant height, seed mass, and phenology did not play any role; however, plant species with higher specific leaf area were more likely to belong to dark species (Fig. 4).

**Fig. 4.**
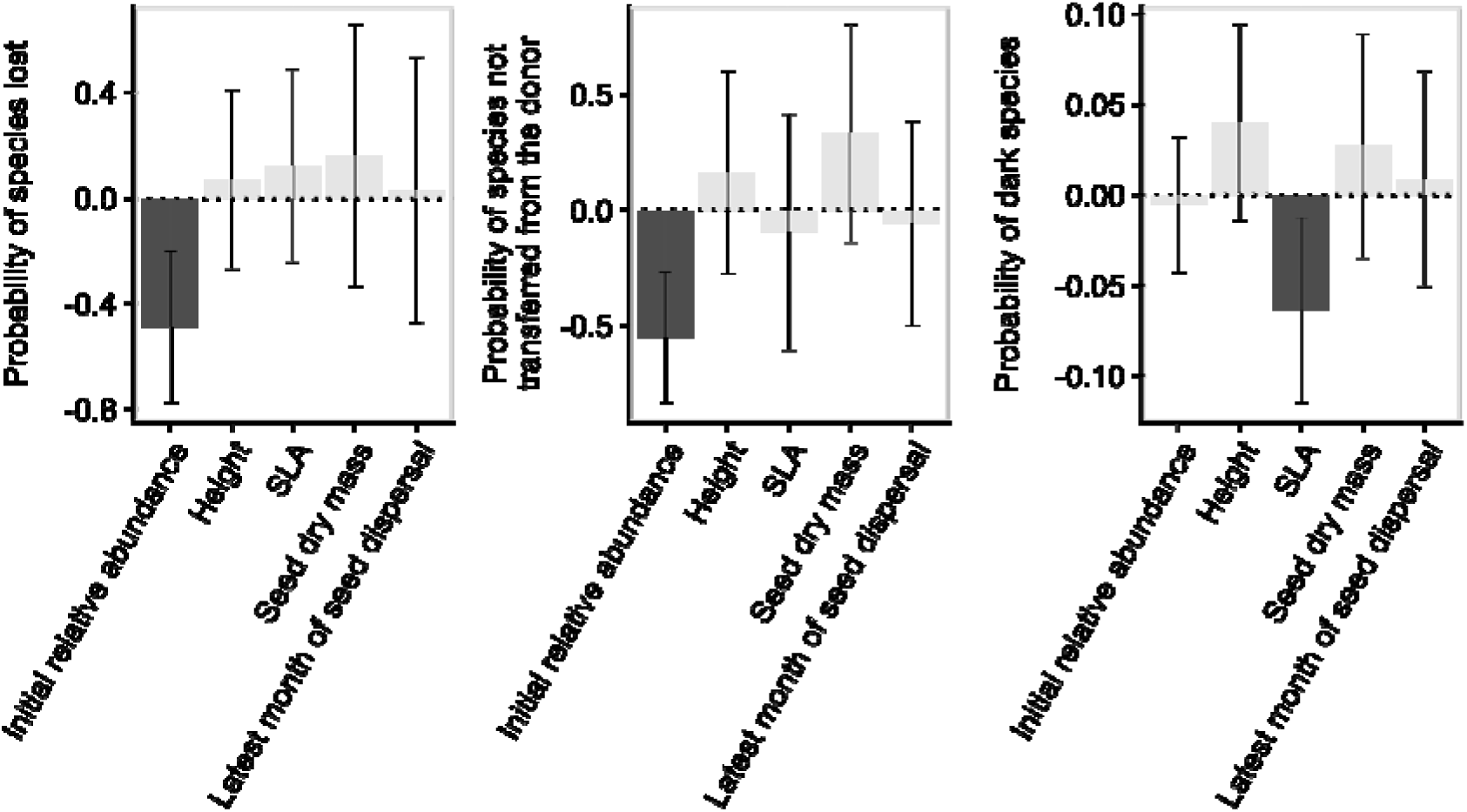
Standardized regression coefficients from multiple regression models showing the effects of initial relative abundance and functional traits on the probability of species lostlost (relative to species persistence), not transferred from the donor (relative to species gained from the donor), and dark diversity in plants. See Table 1 for the definitions of response and predictor variables. Bar plot features as in Fig. 3.

### Invertebrates

In contrast to plants, we recorded higher proportions of species lost in receiver meadows (52–69% for Auchenorrhyncha and 34–48% for Orthoptera). Like for plants, however, we also observed a relatively high proportion of species persisting (47–62% for Auchenorrhyncha and 42–60% for Orthoptera). Yet, more species of invertebrates than plants in the final sampling year originated from species gained from the surroundings (29–48% for Auchenorrhyncha and 36–52% for Orthoptera). Only a small proportion of species were transferred from the donor, with 12–-19% for Auchenorrhyncha and 0-9% for Orthoptera. We found that 83-100% and 79-86% of the unique species of Auchenorrhyncha and Orthoptera were not transferred from the donor to receiver meadows. Moreover, dark diversity accounted for 38–44% of the species not transferred in Auchenorrhyncha and 38–45% in Orthoptera, corresponding to 2.4–2.5 and 0.7–0.8 species, respectively. (Fig. 2 and Fig. S3).

In general, we found weak effects of treatments on overall richness, as well as on the components of species richness change for Auchenorrhyncha and Orthoptera (Fig. 2; Fig. S2; Fig. S3). For Auchenorrhyncha, Hay transfer + deep tillage had the highest proportion of dark diversity, which was significantly higher than hay transfer + shallow tillage. For Orthoptera, the multiplied seeds + deep tillage had the highest proportion of species gained from the surroundings, which was significantly higher than hay transfer + shallow tillage (Fig. 2 and Fig. S2).

Similar to plants, components of species richness change in Auchenorrhyncha were shaped by multiple site characteristics. Higher initial species richness in receiver meadows was associated with a greater proportion of species persisting and a lower proportion of species gained from the surroundings. Similarly, higher initial relative richness was associated with a lower proportion of species not transferred from donors and a lower dark diversity (Fig. 5). In Orthoptera, components of species richness change were largely unrelated to the examined site characteristics, except for a positive relationship between initial species richness in receiver meadows and the proportion of species persisting (Fig. 7). Components of species richness change in both Auchenorrhyncha and Orthoptera were generally not associated with plant biomass or temporal changes in plant diversity metrics, including species richness, Shannon diversity, and Bray–Curtis dissimilarity (Fig. 5; Fig. 7; Fig. S4–S5).

**Fig. 5.**
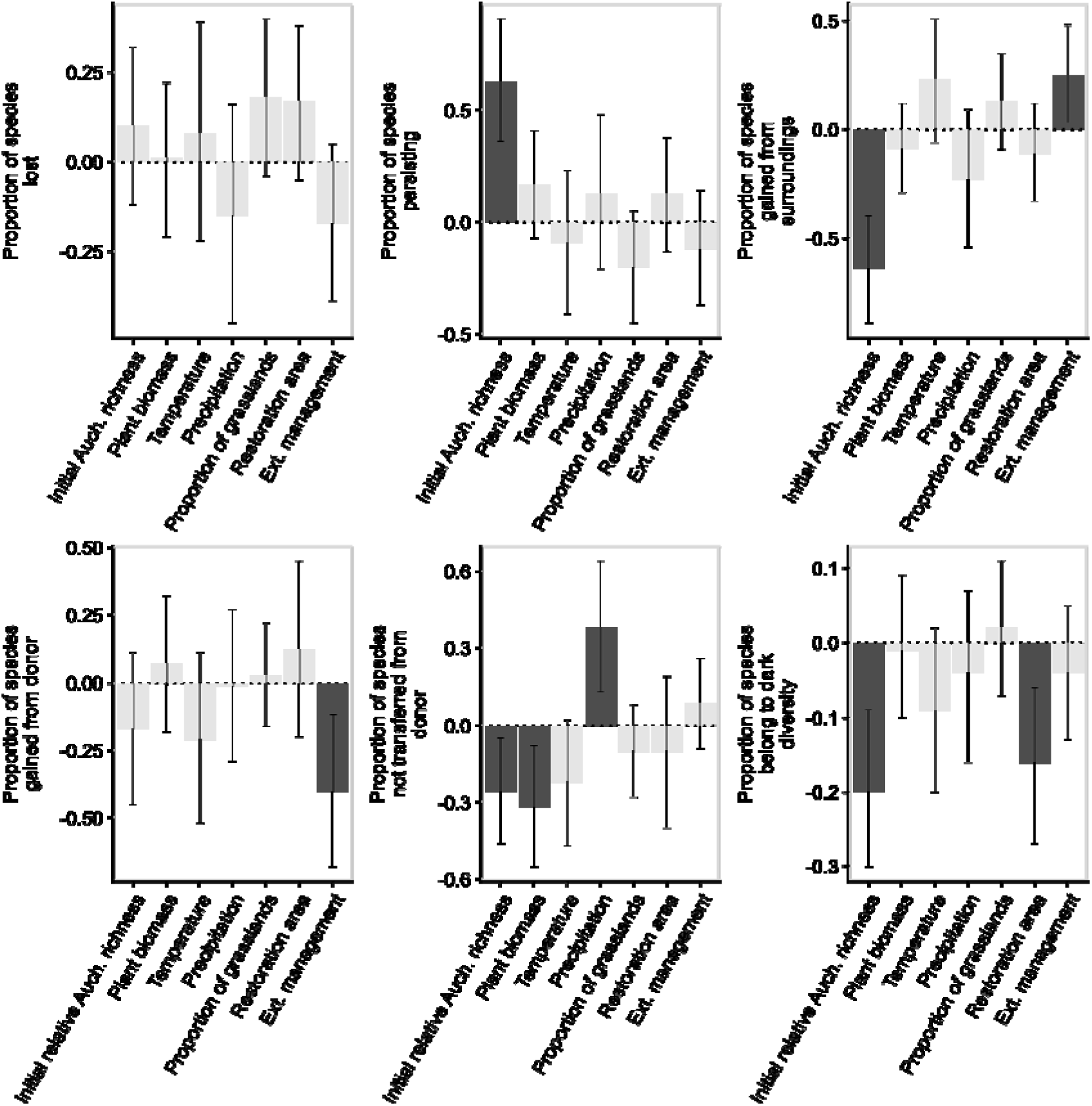
Standardized regression coefficients from multiple regression models showing the effects of community properties, environmental conditions, landscape context, and management history on the components of change of Auchenorrhyncha species richness based on proportional changes. See Table 1 for the definitions of response and predictor variables. Bar plot features as in Fig. 3.

**Fig. 7.**
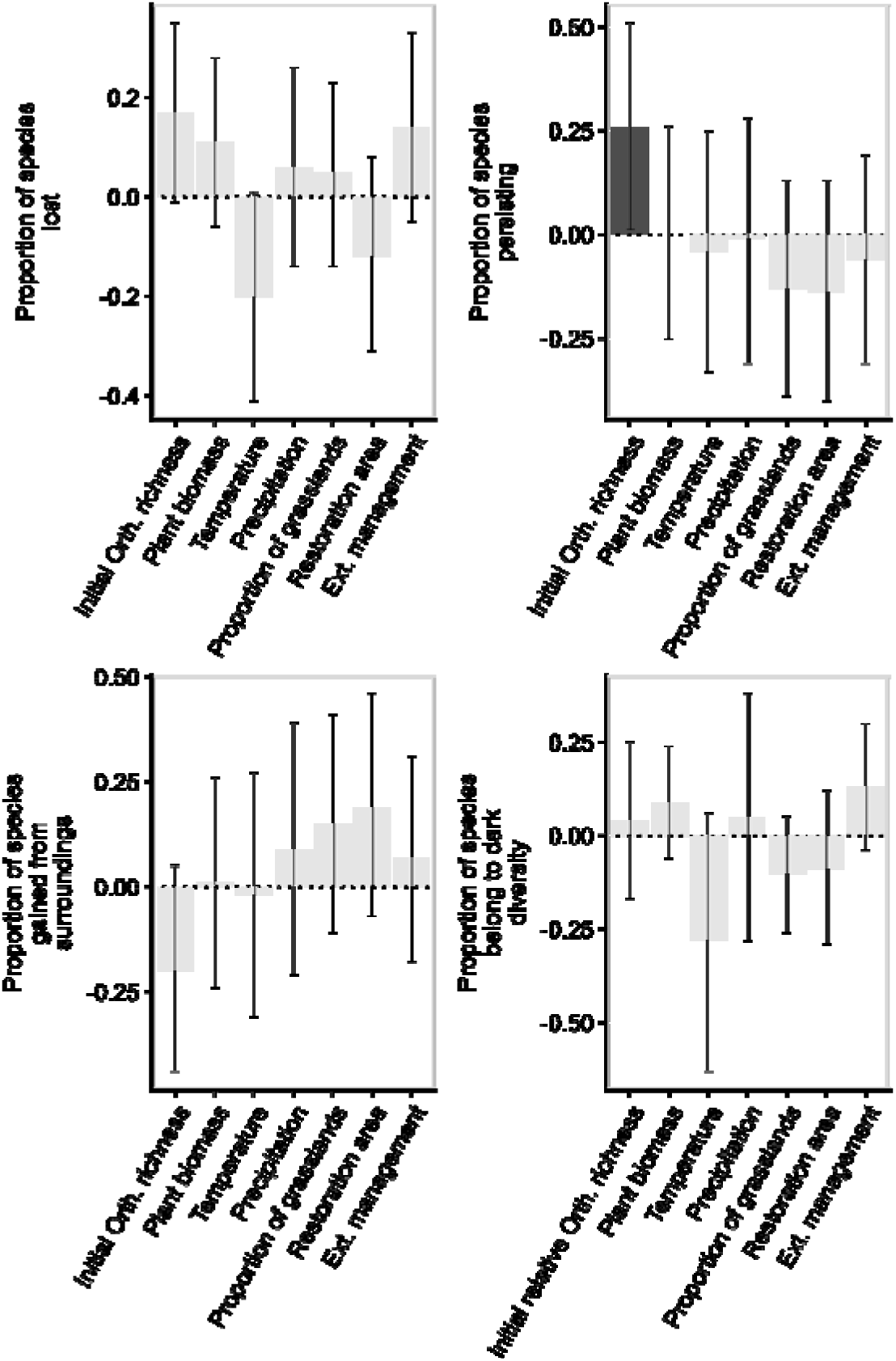
Standardized regression coefficients from multiple regression models showing the effects of community properties, environmental conditions, landscape context, and management history on components of change in Orthoptera species richness based on proportional changes. See Table 1 for the definitions of response and predictor variables. Bar plot features as in Fig. 3.

At the species level, Auchenorrhyncha species with higher relative abundance in receiver meadows showed a lower probability of being lost. Species with higher relative abundance in donor meadows were less likely to belong to dark species (Fig. 6). In contrast, for Orthoptera, no relationships were found between species relative abundance or functional traits and components of species richness change (Fig. 8).

## Discussion

Our study extends previous research in ecological restoration of grasslands by examining the processes underlying species richness change across multiple trophic levels, providing a more mechanistic understanding of community responses to restoration operations. We could evidence that experimental restoration operations altered components of species richness change, with stronger effects in plants than in invertebrates. At the same time, restoration outcomes were strongly shaped by initial species richness in both receiver and donor meadows, highlighting the importance of pre-restoration community properties in determining restoration trajectories. At the species level, we further demonstrate that the probability that a species belongs to the dark species pool depended on its functional traits and/or its initial relative abundance. Finally, we show that components of species richness change in invertebrates were not strongly associated with change in plant communities over time, suggesting a decoupling of responses in plant and invertebrate during restoration.

### Taxon-dependent assembly processes

Although overall species richness did not differ significantly between restoration operations for the three taxonomic groups investigated (Fig. S2), community assembly processes differed markedly between plants and invertebrates. The differences were expressed in the components of species richness change, dark diversity, and the relative importance of management versus site-specific characteristics. Restoration operations did not affect the absolute magnitude of the components of species richness change (Fig. S3) but primarily altered their relative contributions. Additionally, plant community composition change over time differed significantly between direct seeding + deep tillage and either hay transfer + shallow tillage or the control, as indicated by Jaccard and Bray–Curtis dissimilarities. We conclude that the restoration operations primarily affected species composition rather than overall species richness change.

An absence of positive restoration effects on overall plant species richness in our large-scale experiment contrasts with both our initial hypothesis and with former small-scale experiments that have documented successful restoration through hay transfer and/or seed addition (Bischoff et al. 2018; Freitag et al. 2021; Kiehl & Wagner 2006; Orrock et al. 2023; Pywell et al. 2002, 2007; Wagner et al. 2021). A first possible explanation for this discrepancy between small-scale and large-scale experiments is that that plant recruitment and establishment at larger spatial scales are influenced by more complex environmental and soil variables, with environmental factors potentially exerting stronger effects (Rohal et al., 2019). Another possible explanation lies in the relative high initial plant species richness in our receiver meadows at the onset of the experiment (25 ± 4, on average ± SD), which contrasts with previous grassland restoration experiments that were often launched on species-poor agriculture land such as arable land (Laschke et al., 2026). Within the same research framework as here, Forgione et al. (2026) shows a significant increase in plant species richness in all our receiver meadows (+29% on average) between 2018 and 2023. Yet, this amelioration was paralleled by an increase of 16% in the control meadows, suggesting that a mechanism of passive restoration operated in the background, contributing to a large extent to the overall augmentation of plant species richness.

Importantly, the tested restoration operations involving soil disturbance at the onset of our experiment did not reduce invertebrate richness, a welcome result due to the paucity of studies on the response of invertebrates to grassland restoration operations (Slodowicz et al., 2023). This suggests that invertebrates either survived soil disturbance or that sufficient source populations had persisted in the surrounding matrix, e.g. along field margins and/or in adjacent semi-natural habitats, enabling rapid recolonization following vegetation re-establishment (Woodcock et al., 2010; Slodowicz, et al., 2023).

Our results demonstrate that the outcomes of our grassland restoration experiment were shaped by both the operations implemented and the characteristics of the study sites. However, these outcomes are taxon-specific. In plants, the components of species richness change were influenced by the two factors. Overall, restoration operations consistently increased the proportion of species lost and reduced the proportion of species persisting, relative to the control, which may to a large extent reflect a negative impact of the initial soil disturbance. Specifically, hay transfer + deep tillage promoted the establishment of donor-derived species relative to shallow tillage. This effect may reflect stronger suppression of resident vegetation by deeper soil disturbance, thereby reducing interspecific competition and creating more favourable conditions for the establishment of newly introduced species (Davis et al., 2000). We could not evidence differences between the other three restoration methods (hay transfer + deep tillage, direct seeds + deep tillage or multiplied seeds + deep tillage), suggesting that hay transfer + direct seeding perform similarly, in line with recent field-scale restoration experiments conducted in Germany (Laschke et al., 2026).

In contrast to plants, the components of species richness change in Auchenorrhyncha were primarily shaped by site characteristics, while in Orthoptera we could not detect any effect of the restoration method or site characteristics. Stöckli et al. (2021) have established that during restoration operations hay transfer introduced an average of 9.2 living invertebrates per m□². Although Auchenorrhyncha and Orthoptera accounted for only 3.1% by number of the invertebrates transferred, this would contribute to an average total of 1,500 individuals per receiver meadow. Nevertheless, the absence of a difference in the number of recorded invertebrates between receiver meadows with hay transfer and direct seeding (that latter involving no transfer of invertebrates) suggests that direct transfer of invertebrates was ineffective. Instead, colonization from extant populations in the surroundings were likely to contribute to most invertebrates recorded in the last sampling year.

Contrary to previous grassland restoration research that often accounts for just a few site characteristics, the present study evaluated the relative importance of a wide array of characteristics — ecological community properties, environmental conditions, landscape context, and management history — in shaping the processes underlying changes in species richness. We demonstrate here, first, that initial conditions, in both receiver and donor meadows, are key to predict processes driving species richness change during restoration operations for plants and Auchenorrhyncha. Second, local soil nutrient is also important: sites with higher soil nitrogen had a greater proportion of species persisting maybe due to the strong dominance of nitrogen-adapted plants that outcompete exogenous propagules, a well-known ecological phenomenon (Freitag et al. 2021). Soil pH also strongly influenced several components of change in plant species richness, suggesting that abiotic filtering shapes the processes underlying species richness change. On the contrary, Orthoptera showed only weak relationships with all site characteristics. On the one hand, this taxon consists mostly of ubiquitous generalists (Devictor et al., 2010; Joern, 1979); on the other hand, the local community is not very speciose, with a maximum of seven species recorded in a meadow, which provides limited power for statistical analyses.

### Dark diversity and limits to community assembly

The patterns of dark diversity we evidenced revealed substantial unrealized potential among the three investigated taxa: many species potentially suitable to site conditions were missing in the receiver meadows. For plants, the multiplied seeds + deep tillage introduced a high number of unique species into the receiver community but this treatment yielded the second highest proportion of species not transferred from the donor, and yet the lowest dark diversity among the treatments. This indicates that many species of the commercial seed mixture we used had a weak realised and potential establishment under the local environmental circumstances. Concerning Auchenorrhyncha, the negative association between the initial relative species richness and dark diversity suggests that communities having more similar number of species to donor richness contained fewer dark species and therefore may reduce potential for further colonisation. Dark diversity in Auchenorrhyncha was additionally related to pre-seeding soil preparation and area of the receiver meadow, highlighting the importance of both restoration operations and landscape context in shaping dark diversity. These results demonstrate that incorporating dark diversity provides insights into restoration potential beyond changes in species richness alone.

### Species-level mechanisms and assembly filters

Ecological research has shown that initial species abundance and functional traits mediate plant community responses during the species assembly process (Suding et al., 2005). Yet, their effects on the components of species richness change across trophic levels have so far not been assessed simultaneously in any grassland restoration experiment. The present study fills this gap in knowledge, demonstrating how initial species relative abundance and species traits jointly shape restoration outcomes. Our species-level analyses established the existence of trait- and abundance-based assembly filters, which are both taxon-specific. However, there was limited consistency in species-level patterns across study sites, particularly for plants, with dark species varying markedly among restoration operations and sites, to the extent that we could not identify a set of common species across our study sites (Table S2-Table S4). Functional traits traditionally predicting which species are top candidates for dark diversity (i.e. low dispersal capacity, low competitive ability and high stress-tolerance; Belinchón et al., 2020) proved ineffective in our experiment. Nevertheless, in plants and Auchenorrhyncha, species with higher initial abundance were less likely to be lost or not transferred from donor-derived assemblages, corroborating community assembly theory (Suding et al., 2005). Moreover, higher specific leaf area, potentially associated with fast-growing strategies, increased the likelihood that a given plant species belongs to dark diversity. In contrast, no trait effects or effects of initial abundance were detected on Orthoptera, indicating fundamentally different assembly filters in this taxon.

### Decoupling between vegetation and invertebrates

The observation that invertebrate responses were not associated with plant biomass or changes in plant diversity over time suggests that plant-based metrics are insufficient proxies for predicting components of species richness change in invertebrates. This result contradicts the widespread view that the restoration of invertebrate communities chiefly depends on the restoration of a rich plant community (e.g., Alignan et al. 2018; Alison et al. 2017; Neff et al. 2020; Woodcock et al. 2010). Our results on Orthoptera furthermore substantiate the claim that invertebrate restoration may primarily be regulated by environmental factors, and not by the seeding method (Kiehl & Wagner 2006), notably by farming conditions in the wider landscape (Seibold et al., 2019). This means that targeted translocations of live invertebrates might be better options for increasing species richness at receiver meadows (Knop et al., 2008; Seibold et al., 2019). The present results also emphasize that a multi-taxon approach is needed to properly evaluate grassland restoration success and frame general recommendations.

### Conclusion and management implications

Our large-scale, controlled experiment shows that the four different restoration operations tested appear fairly equivalent in terms of restoration power. None altered overall species richness across the three investigated taxa, although they substantially shaped the underlying community assembly process. Also, pre-restoration community properties (e.g. species richness and relative species abundance) played important roles in determining restoration trajectories. This bears three key implications for grassland restoration research and practice. First, restoration should prioritize plant species that are strongly associated with dark diversity. Planning seed collection targeting dark plant diversity and relying on seed mixtures in which dark species are over-represented may boost the establishment of a rich ecological community. Second, restoration success must be evaluated using multi-trophic assessments rather than focusing exclusively on vegetation in order to avoid overestimating the recovery of invertebrates. Third, integrating broader environmental conditions into restoration strategies is critical for promoting invertebrate biodiversity.

## Supporting information

Supplementary file for Dark diversity reveals assembly constraints across trophic levels in large-scale grassland restoration

## AUTHOR CONTRIBUTIONS

Jean-Yves Humbert and Raphaël Arlettaz conceived the project. Jean-Yves Humbert,Laura Forgione, Michael Wyss, Daniel Slodowicz, and Gino Enz collected the data. Qingqing Chen and Jean-Yves Humbert developed the research framework; Qingqing Chen analyzed the data; Qingqing Chen led the writing of the manuscript. All authors contributed critically to the drafts and gave final approval for publication.

## ACKNOWLEDGEMENTS

We thank all involved farmers and cantons (i.e. Bern, Fribourg, Luzern, Solothurn and Vaud) for their collaboration. We thank the stakeholder accompanying group for its valuable advice and logistic support throughout the project. We thank Lara Schärer, Astrance Fenestraz, Roman Roth, Sarah Ettlin, and Yasemin Kurtogullari for field assistance.

## FUNDING INFORMATION

Swiss National Science Foundation (grant no. 310030_215272 to Raphaël Arlettaz and Jean-Yves Humbert) and the Federal Office for Agriculture.

## CONFLICT OF INTEREST STATEMENT

Authors declare that they have no competing interests.

---

**Fig. 6.** Standardized regression coefficients from multiple regression models for the effects of relative abundance and functional traits on probability of species lost (relative to species persistence), species not transferred from the donor (relative to species gained from the donor), and dark species for Auchenorrhyncha. See Table 1 for the definitions of response and predictor variables. Bar plot features as in Fig. 3.

**Fig. 8.** Standardized regression coefficients from multiple regression models for the effects of relative abundance and functional traits on probability of species lost (relative to species persistence), species not transferred from the donor (relative to species gained from the donor), and dark species for Orthoptera. See Table 1 for the definitions of response and predictor variables. Bar plot features as in Fig. 3.

## Notes

### Competing Interest Statement

The authors have declared no competing interest.

