## Supplementary file for Dark diversity reveals assembly constraints across trophic levels in large-scale grassland restoration for "Dark diversity reveals assembly constraints across trophic levels in large-scale grassland restoration"

11 September 2026

The file contains:

Figure S1–Figure S5

Table S1–Table S4

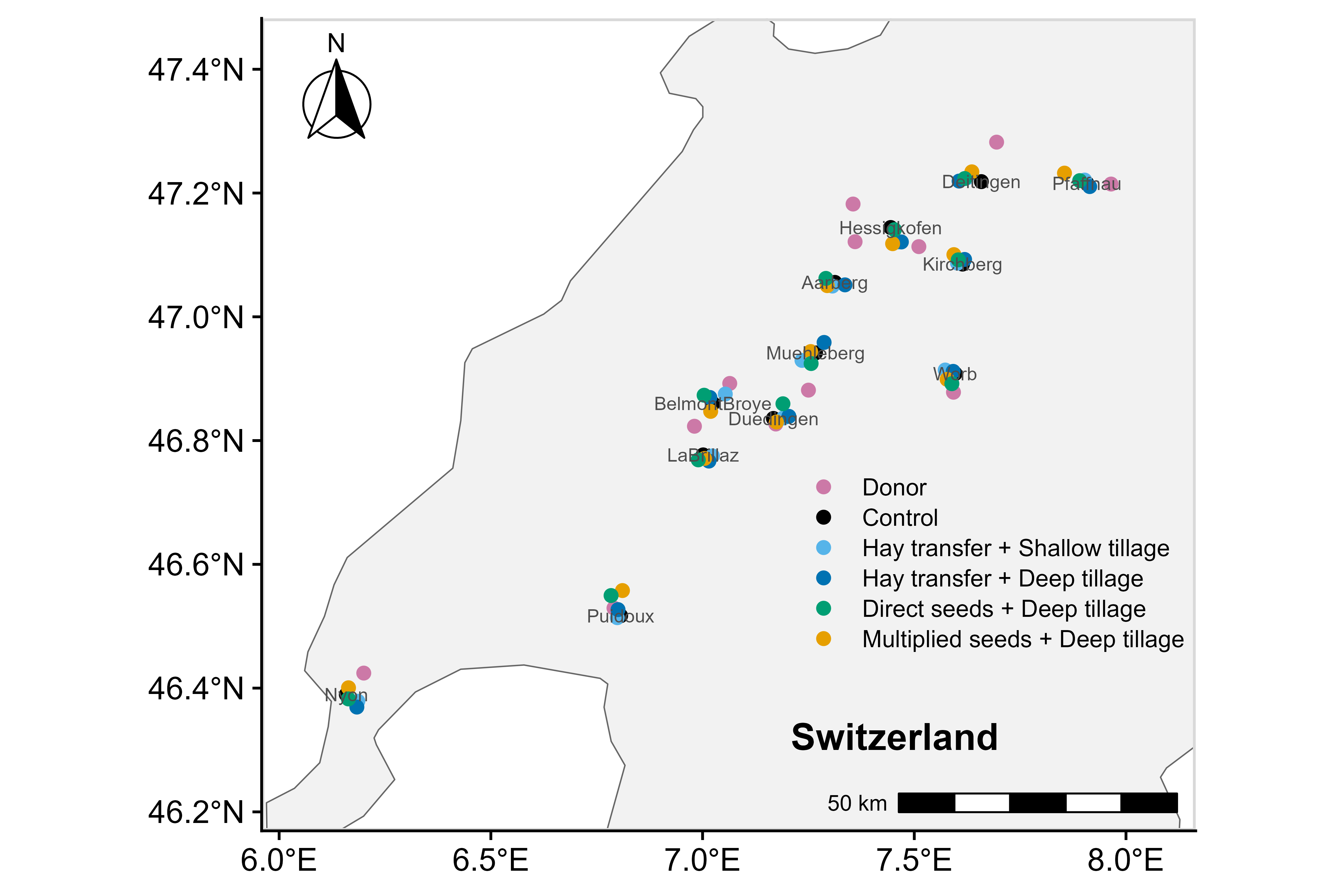

Fig. S1. Geolocation of the study sites.

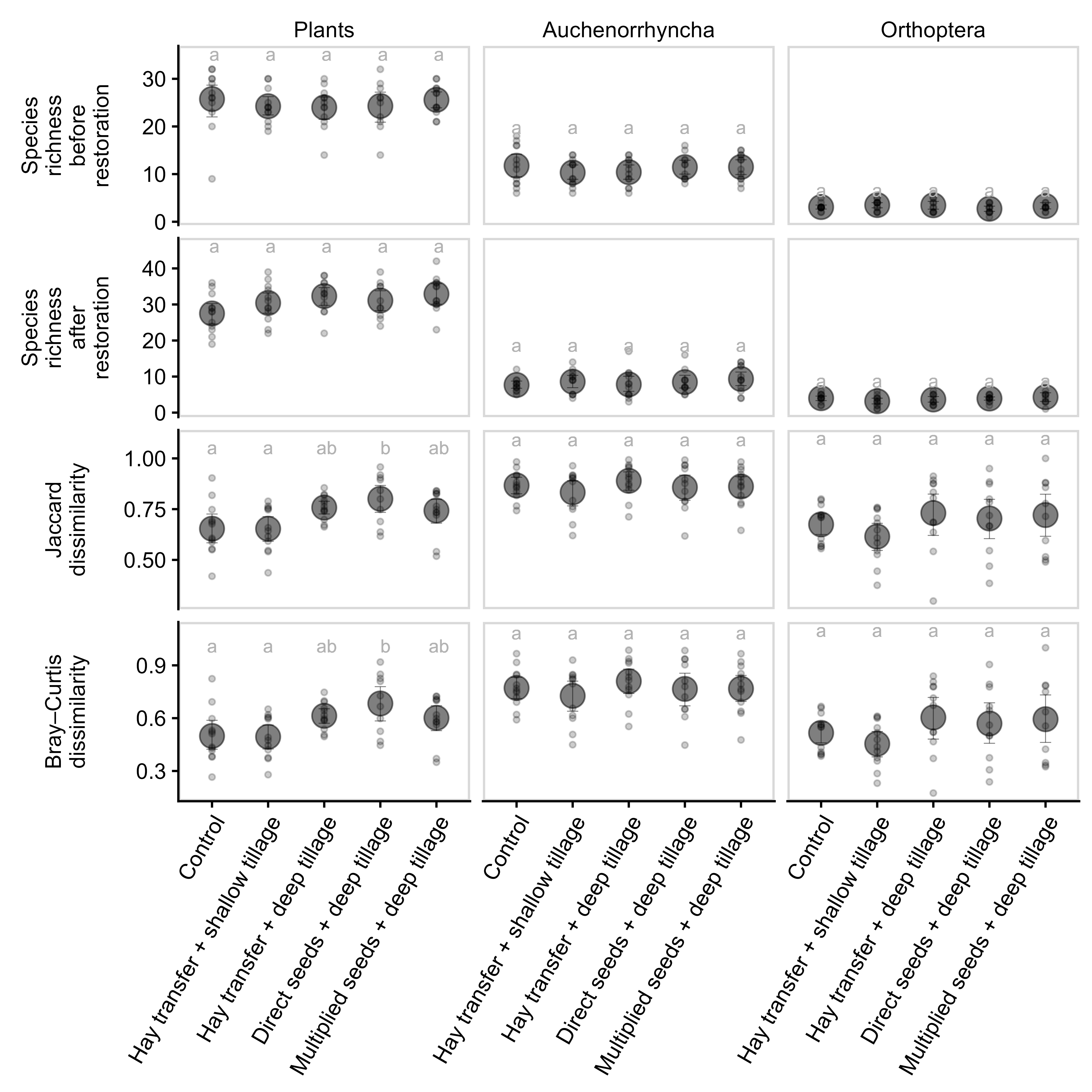

Fig. S2. Treatment effects on species richness and community dissimilarity for plants, Auchenorrhyncha, and Orthoptera before restoration (in 2018) and in the last sampling year. Species richness for plants, Auchenorrhyncha, and Orthoptera was recorded within 16 m², 0.8 m², and 16 m², respectively. Small points represent region-level raw data. Large points and error bars represent means and 95% bootstrapped confidence intervals. Different letters indicate significant differences at α = 0.05 with Tukey-adjusted multiple comparisons.

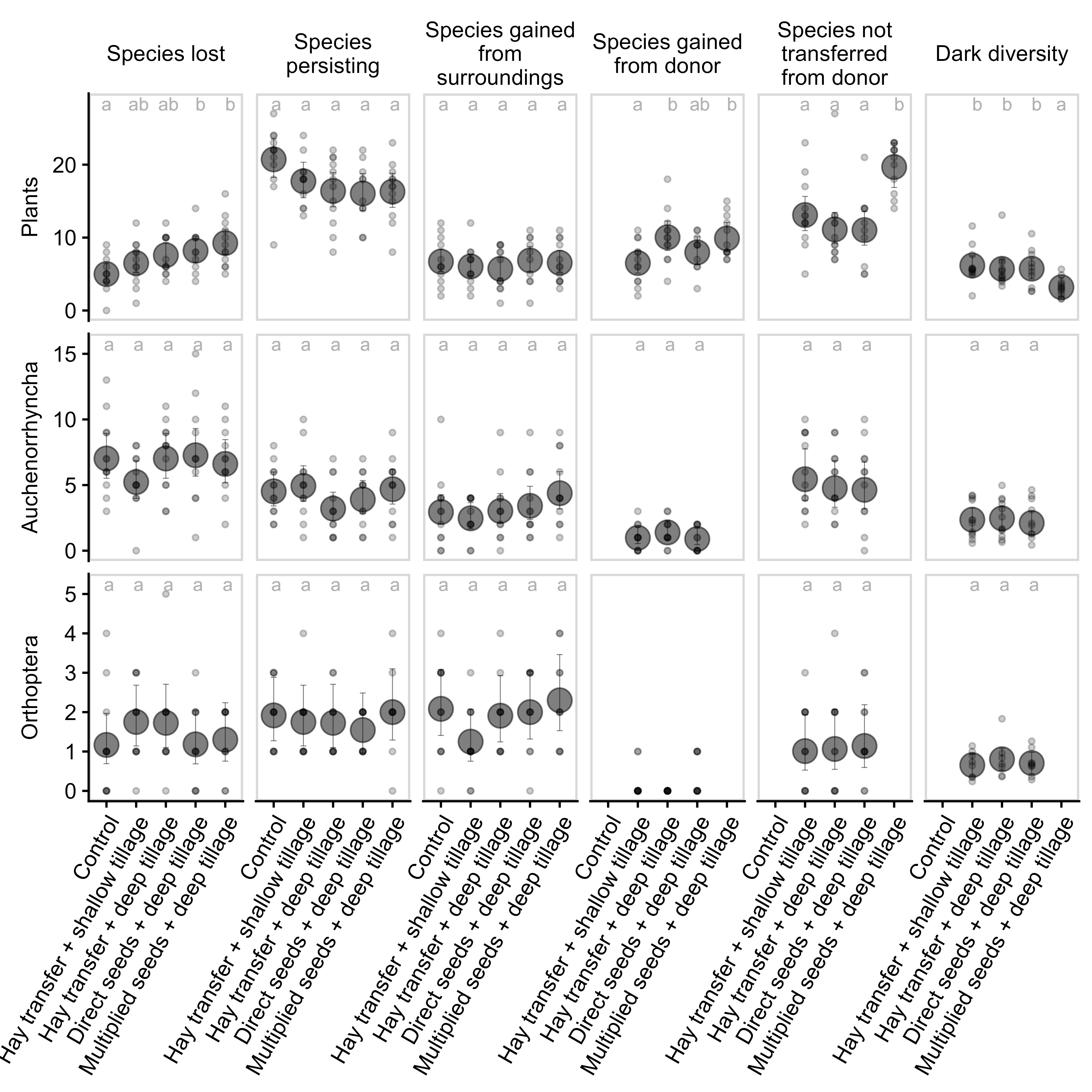

Fig. S3. Treatment effects on components of species richness change (absolute change) for plants, Auchenorrhyncha, and Orthoptera. Components defined in Fig. 1. Small points show region-level raw data; triangles indicate observations excluded from the models. Large points and error bars represent estimated means and 95% confidence intervals. Different letters indicate significant differences at α = 0.05 with Tukey-adjusted multiple comparisons.

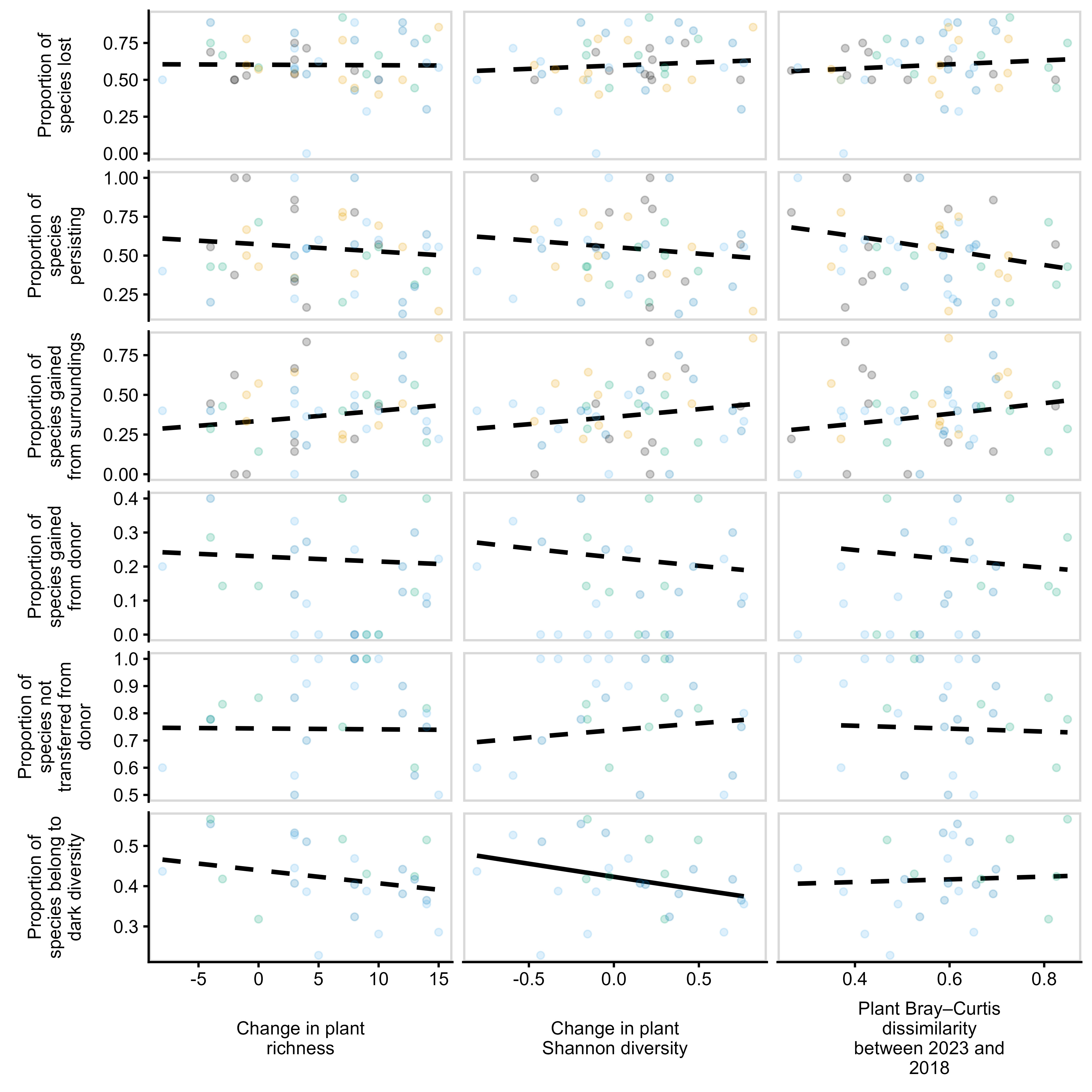

**Fig. S4.** Predicted relationships between the change in plant diversity from 2018 to 2023 and each component of Auchenorrhyncha species richness change. Solid lines indicate statistically significant relationships, whereas dashed lines indicate non-significant relationships.

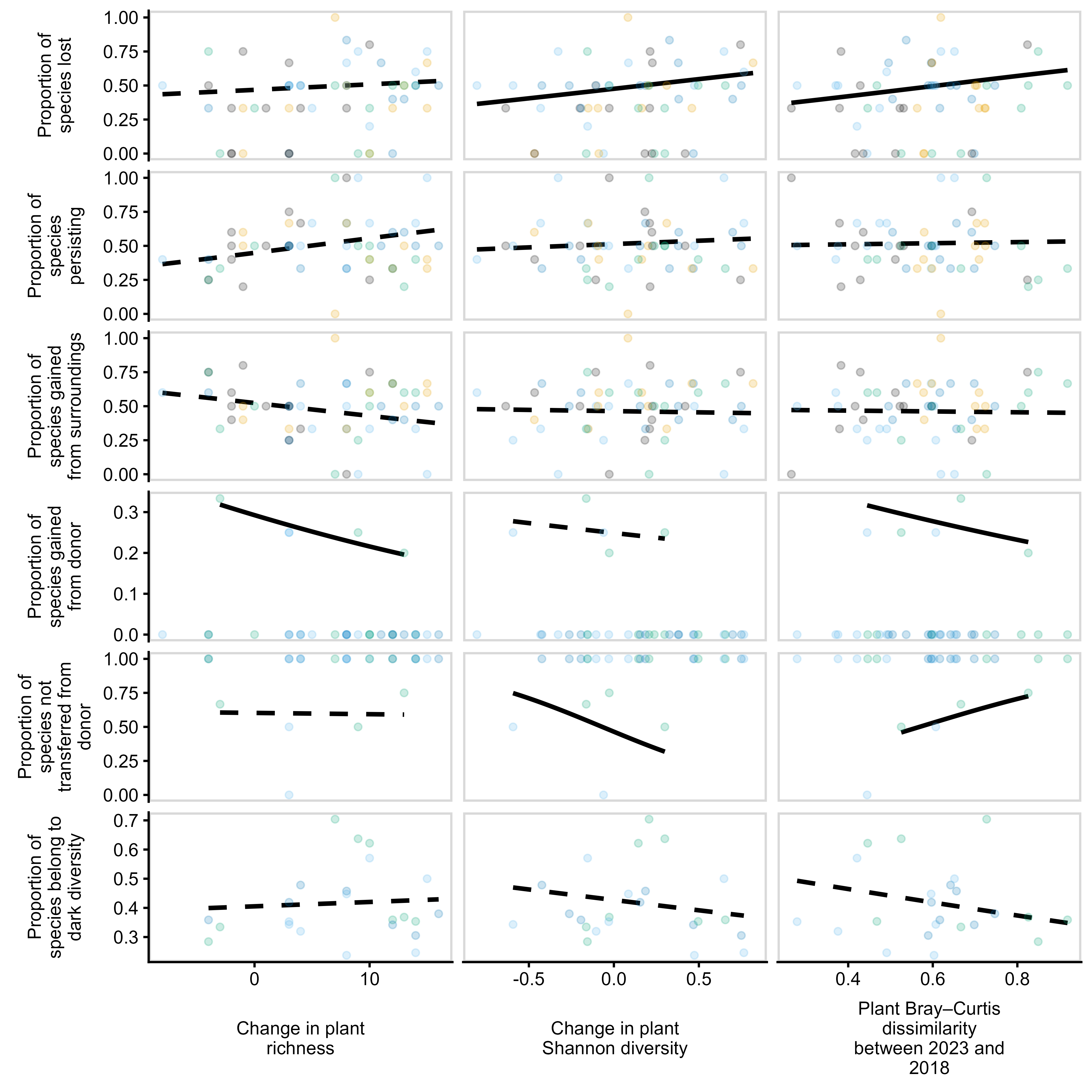

**Fig. S5.** Predicted relationships between the change in plant diversity from 2018 to 2023 and each component of Orthoptera species richness change. Solid lines indicate statistically significant relationships, whereas dashed lines indicate non-significant relationships.

Table S1. Summary of the average number of shared and unique species recorded in donor meadows for each restoration treatment. DS-DT: Direct seeds + deep tillage; HT-DT: Hay transfer + deep tillage; HT-ST: Hay transfer + shallow tillage; MS-DT: Multiplied seeds + deep tillage

| Taxa groups | Treatment | Total number of species in donor | Shared species between donor and receiver meadows | Number of unique species | Proportion of unique species |
| --- | --- | --- | --- | --- | --- |
| Auchenorrhyncha | DS-DT | 11 | 5 | 6 | 0.49 |
| Auchenorrhyncha | HT-DT | 11 | 4 | 7 | 0.58 |
| Auchenorrhyncha | HT-ST | 11 | 4 | 7 | 0.62 |
| Orthoptera | DS-DT | 4 | 2 | 2 | 0.40 |
| Orthoptera | HT-DT | 4 | 3 | 1 | 0.29 |
| Orthoptera | HT-ST | 4 | 2 | 1 | 0.36 |
| Plants | DS-DT | 34 | 15 | 19 | 0.55 |
| Plants | HT-DT | 34 | 13 | 21 | 0.61 |
| Plants | HT-ST | 34 | 15 | 20 | 0.58 |
| Plants | MS-DT | 38 | 8 | 30 | 0.79 |

Table S2. Summary of dark species in plants. Shown are the top five species exhibiting the highest and lowest probabilities of being classified as dark species. For species classified as dark species in more than one region, the reported probability is the mean probability across regions. DS-DT: Direct seeds + deep tillage; HT-DT: Hay transfer + deep tillage; HT-ST: Hay transfer + shallow tillage; MS-DT: Multiplied seeds + deep tillage

| Treatment | Standard taxon | Number of regions species not transferred and recorded as potential dark species | Probability of dark species |
| --- | --- | --- | --- |
| DS-DT | *Leontodon hispidus* | 1 | 0.73 |
| DS-DT | *Briza media* | 2 | 0.70 |
| DS-DT | *Leucanthemum vulgare* | 1 | 0.68 |
| DS-DT | *Helictotrichon pubescens* | 3 | 0.63 |
| DS-DT | *Medicago sativa* | 2 | 0.61 |
| DS-DT | *Veronica chamaedrys* | 3 | 0.36 |
| DS-DT | *Veronica arvensis* | 1 | 0.34 |
| DS-DT | *Ranunculus acris* | 1 | 0.28 |
| DS-DT | *Cerastium fontanum* | 1 | 0.26 |
| DS-DT | *Taraxacum spec* | 1 | 0.23 |
| HT-DT | *Leontodon hispidus* | 1 | 0.71 |
| HT-DT | *Anthriscus sylvestris* | 1 | 0.69 |
| HT-DT | *Colchicum autumnale* | 1 | 0.69 |
| HT-DT | *Knautia arvensis* | 1 | 0.67 |
| HT-DT | *Centaurea scabiosa* | 1 | 0.66 |
| HT-DT | *Lolium multiflorum* | 1 | 0.32 |
| HT-DT | *Veronica chamaedrys* | 1 | 0.32 |
| HT-DT | *Veronica arvensis* | 1 | 0.31 |
| HT-DT | *Festuca rubra* | 2 | 0.31 |
| HT-DT | *Heracleum sphondylium* | 1 | 0.29 |
| HT-ST | *Carex hirta* | 1 | 0.61 |
| HT-ST | *Carex panicea* | 1 | 0.61 |
| HT-ST | *Lactuca serriola* | 1 | 0.61 |
| HT-ST | *Scabiosa columbaria* | 1 | 0.61 |
| HT-ST | *Veronica filiformis* | 1 | 0.61 |
| HT-ST | *Vicia sepium* | 1 | 0.37 |
| HT-ST | *Veronica chamaedrys* | 2 | 0.29 |
| HT-ST | *Dactylis glomerata* | 1 | 0.29 |
| HT-ST | *Veronica arvensis* | 2 | 0.27 |
| HT-ST | *Trifolium repens* | 1 | 0.27 |
| MS-DT | *Picris hieracioides* | 1 | 0.68 |
| MS-DT | *Lathyrus pratensis* | 1 | 0.57 |
| MS-DT | *Campanula patula* | 3 | 0.56 |
| MS-DT | *Salvia pratensis* | 8 | 0.55 |
| MS-DT | *Clinopodium vulgare* | 2 | 0.53 |
| MS-DT | *Leucanthemum vulgare* | 1 | 0.31 |
| MS-DT | *Trisetum flavescens* | 2 | 0.29 |
| MS-DT | *Arrhenatherum elatius* | 1 | 0.26 |
| MS-DT | *Plantago lanceolata* | 1 | 0.24 |
| MS-DT | *Crepis biennis* | 1 | 0.21 |

Table S3. Summary of dark species in Auchenorrhyncha. Shown are the top five species exhibiting the highest and lowest probabilities of being classified as dark species. For species classified as dark species in more than one region, the reported probability is the mean probability across regions. DS-DT: Direct seeds + deep tillage; HT-DT: Hay transfer + deep tillage; HT-ST: Hay transfer + shallow tillage; MS-DT: Multiplied seeds + deep tillage

| Treatment | Standard taxon | Number of regions species not transferred and recorded as potential dark species | Probability of dark species |
| --- | --- | --- | --- |
| DS-DT | *Tettigometra impressopunctata* | 1 | 0.67 |
| DS-DT | *Arocephalus longiceps* | 1 | 0.64 |
| DS-DT | *Philaenus spumarius* | 1 | 0.58 |
| DS-DT | *Ribautodelphax imitans* | 2 | 0.56 |
| DS-DT | *Graphocraerus ventralis* | 1 | 0.55 |
| DS-DT | *Laodelphax striatella* | 1 | 0.23 |
| DS-DT | *Forcipata citrinella* | 1 | 0.20 |
| DS-DT | *Unidentified cicadomorpha* | 1 | 0.20 |
| DS-DT | *Acanthodelphax spinosa* | 1 | 0.18 |
| DS-DT | *Xanthodelphax stramineus* | 1 | 0.14 |
| HT-DT | *Javesella pellucida* | 1 | 0.64 |
| HT-DT | *Tettigometra impressopunctata* | 1 | 0.62 |
| HT-DT | *Ribautodelphax imitans* | 2 | 0.58 |
| HT-DT | *Errastunus ocellaris* | 2 | 0.57 |
| HT-DT | *Euscelis incisus* | 1 | 0.57 |
| HT-DT | *Xanthodelphax stramineus* | 2 | 0.38 |
| HT-DT | *Eupteryx notata* | 2 | 0.36 |
| HT-DT | *Anoscopus serratulae* | 1 | 0.28 |
| HT-DT | *Graphocraerus ventralis* | 1 | 0.26 |
| HT-DT | *Anaceratagallia ribauti* | 2 | 0.24 |
| HT-ST | *Philaenus spumarius* | 2 | 0.61 |
| HT-ST | *Tettigometra impressopunctata* | 1 | 0.61 |
| HT-ST | *Megophtalmus scanicus* | 1 | 0.55 |
| HT-ST | *Ribautodelphax imitans* | 2 | 0.52 |
| HT-ST | *Euscelis incisus* | 1 | 0.51 |
| HT-ST | *Anaceratagallia ribauti* | 1 | 0.24 |
| HT-ST | *Empoasca pteridis* | 1 | 0.21 |
| HT-ST | *Jassargus obtusivalvis* | 1 | 0.21 |
| HT-ST | *Psammotettix confinis* | 1 | 0.18 |
| HT-ST | *Javesella pellucida* | 1 | 0.18 |

Table S4. Summary of dark species in Orthoptera. Shown are the top five species exhibiting the highest and lowest probabilities of being classified as dark species. For species classified as dark species in more than one region, the reported probability is the mean probability across regions. DS-DT: Direct seeds + deep tillage; HT-DT: Hay transfer + deep tillage; HT-ST: Hay transfer + shallow tillage; MS-DT: Multiplied seeds + deep tillage

| Treatment | Standard taxon | Number of regions species not transferred and recorded as potential dark species | Probability of dark species |
| --- | --- | --- | --- |
| DS-DT | *Roeseliana roeselii* | 2 | 0.53 |
| DS-DT | *Chorthippus brunneus* | 2 | 0.47 |
| DS-DT | *Leptophyes punctatissima* | 1 | 0.44 |
| DS-DT | *Phaneroptera falcata* | 1 | 0.44 |
| DS-DT | *Metrioptera parapleurus* | 4 | 0.42 |
| DS-DT | *Bicoloria bicolor* | 1 | 0.41 |
| DS-DT | *Stauroderus scalaris* | 1 | 0.41 |
| DS-DT | *Stenobothrus lineatus* | 2 | 0.33 |
| DS-DT | *Gomphocerippus rufus* | 1 | 0.29 |
| HT-DT | *Phaneroptera falcata* | 2 | 0.50 |
| HT-DT | *Leptophyes punctatissima* | 1 | 0.44 |
| HT-DT | *Bicoloria bicolor* | 1 | 0.42 |
| HT-DT | *Stauroderus scalaris* | 1 | 0.42 |
| HT-DT | *Roeseliana roeselii* | 4 | 0.40 |
| HT-DT | *Metrioptera parapleurus* | 2 | 0.38 |
| HT-DT | *Stenobothrus lineatus* | 2 | 0.36 |
| HT-DT | *Chorthippus brunneus* | 1 | 0.31 |
| HT-ST | *Leptophyes punctatissima* | 1 | 0.57 |
| HT-ST | *Phaneroptera falcata* | 2 | 0.50 |
| HT-ST | *Chorthippus biguttulus* | 2 | 0.42 |
| HT-ST | *Stenobothrus lineatus* | 2 | 0.35 |
| HT-ST | *Roeseliana roeselii* | 2 | 0.35 |
| HT-ST | *Bicoloria bicolor* | 1 | 0.31 |
| HT-ST | *Stauroderus scalaris* | 1 | 0.31 |
| HT-ST | *Chorthippus brunneus* | 3 | 0.30 |
